# KLF7 is a general inducer of human pluripotency

**DOI:** 10.1101/2023.09.06.556189

**Authors:** Mattia Arboit, Irene Zorzan, Marco Pellegrini, Paolo Martini, Elena Carbognin, Graziano Martello

## Abstract

Pluripotency is the capacity to give rise to all differentiated cells of the body and the germ line and is governed by a self-reinforcing network of transcription factors. The forced expression of only some of these factors enables the reprogramming of somatic cells to pluripotency. In murine cells, several kruppel-like factors (KLFs) have been identified as stabilisers and inducers of pluripotency. Human somatic cells are routinely reprogrammed by expression of KLF4 in combination with OCT4, SOX2 and cMYC (OSKM). An extensive transcriptome analysis revealed, however, that KLF4 is barely expressed in conventional human pluripotent stem cells (PSCs). Here we show that KLF7 is robustly expressed in conventional human PSCs and it allows transcription factor-mediated somatic reprogramming. Moreover, we find that KLF7 is highly expressed in naive PSCs and its forced expression in conventional hPSCs induces upregulation of naive markers and boosts efficiency of chemical resetting to naive PSCs, overall suggesting that KLF7 is a general human pluripotency factor and an inducer of pluripotency.

## INTRODUCTION

Pluripotent stem cells have been originally derived by culturing epiblast cells of early mouse embryos as mESCs^1,2^. The molecular characterisation of these cells allowed the identification of specific pluripotency factors which enabled the induction of pluripotency from somatic cells by transcription factor-mediated reprogramming, and the derivation of induced Pluripotent Stem cells (iPSCs)^3^.

To generate iPSCs, the forced expression of a cocktail of four factors (OCT4, SOX2, KLF4 and cMYC) (OSKM) was firstly employed for both murine and human somatic reprogramming^3,4^, despite the species-specific differences.

Human conventional PSCs, derived from human blastocysts as human embryonic stem cells (hESCs)^5^ or generated by reprogramming of somatic cells^6^ as iPSCs, resemble a developmental stage called primed pluripotency corresponding to the epiblast of the post-implantation blastocyst^7,8^. Recently, human naive pluripotent cells, mirroring a second pluripotent state resembling the pre-implantation embryo, have been derived directly from the human pre-implantation embryo^9^, by reprogramming of fibroblasts^10–13^, by transgene-mediated resetting of human conventional PSCs^14,15^ and by chemical resetting^15–17^.

In the murine system, several Kruppel-like factors (Klf2/4/5) are important for the maintenance of pluripotency in naive cells and are absent in primed EpiSCs^18^. Indeed, these Kruppel-like factors have been used to generate naive iPS cells from mouse fibroblasts, in combination with OSM, and are all able to reset EpiSCs to naive pluripotency^19–21^.

In human naive cells, KLF4, KLF5 and KLF17 are highly expressed^22–24^, but their expression in conventional/primed PSCs remains unexplored. Analysis of human pre- and post-implantation epiblast cells indicate that these Kruppel-like factors are expressed specifically in the naive state^25^. Indeed, KLF17 and KLF4 have been shown to be a powerful inducer of naive pluripotency *in vitro*^26^. Surprisingly, however, KLF4 is also routinely used to generate conventional human iPSCs.

We performed a thorough transcriptional analysis of conventional PSCs cultured under different conditions and consistently failed to detect robust KLF4 expression. In contrast, a human-specific KLF, named KLF7, is robustly expressed in conventional PSCs and supports pluripotency downstream of TGF-beta^27^. Expression of KLF7 alongside reprogramming factors OCT4, SOX2 and cMYC, enables robust human somatic reprogramming. KLF7 is highly expressed also in human naive PSCs, and its forced expression in conventional PSCs maximises chemical resetting to naive pluripotency. Thus, KLF7 is a general inducer of human pluripotency.

## RESULTS

### KLF7 enables induction of pluripotency

Induction of pluripotency in human somatic cells can be achieved via the expression of different combinations of the factors OCT4, SOX2, KLF4, MYC, NANOG, LIN28A (OSKM and OSNL), which leads to the generation of induced Pluripotent Stem Cells (iPSCs)^4,6^. We verified the absolute expression of such reprogramming factors in several human PSC and iPSC lines by interrogating previously published data^10,12,27–33^ and observed that expression of the pluripotency factor KLF4 was barely detectable (Fig. 1a). This prompted us to measure the expression of other Kruppel-like factors previously involved in pluripotency maintenance or induction in either human or murine cells (KLF2/4/5/7/17) and noticed that KLF7 is the most represented amongst others, at levels comparable to pluripotency markers such as PRDM14 and ZNF398^27^ (Fig. 1b). The other KLFs analysed were all expressed at low levels, comparable to markers of early differentiation such as SOX1 and TBXT1 (also known as BRACHYURY or T). Of note, we systematically failed to detect expression of KLF4, regardless of the origin of PSCs analysed (embryo-derived (ESCs) or induced (iPSCs)) (Fig. 1c).

**Fig. 1:**
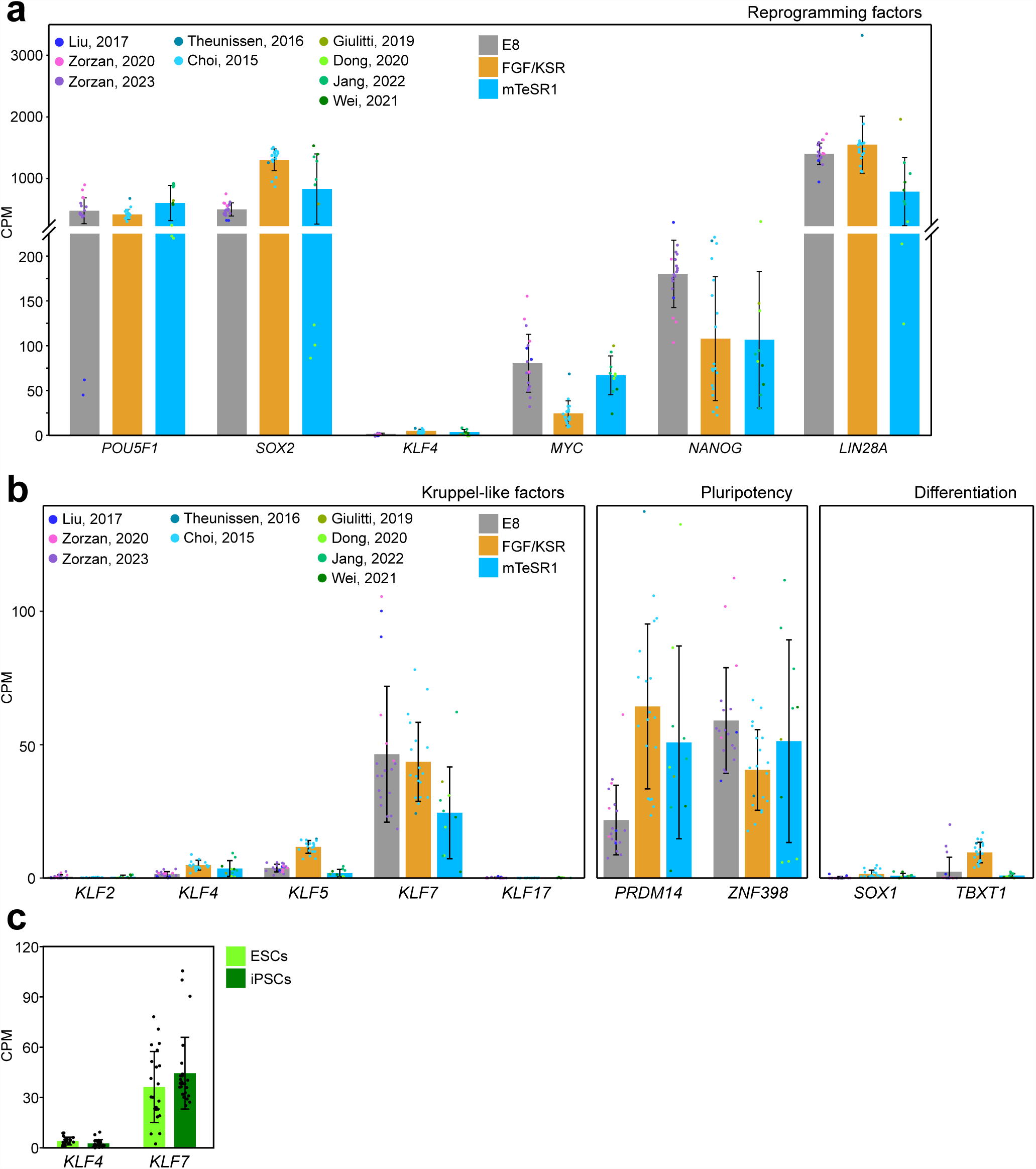
Expression of reprogramming factors and KLFs in conventional human PSCs. a: Barplot showing the absolute expression, measured by RNAseq, of reprogramming factors in conventional hPSCs cultures in 3 distinct culture conditions. Data were obtained from^10,12,27–33^. Mean +/- SD of at least 8 biological replicates is shown. b: Barplot showing the absolute expression, measured by RNAseq, of Kruppel-like factors, pluripotency and differentiation genes in conventional hPSCs, cultured in 3 distinct culture conditions. Data were obtained from^10,12,27–33^. Mean +/- SD of at least 8 biological replicates is shown. c: Barplot showing the absolute expression, measured by RNAseq, of Kruppel-like factors KLF4 and KLF7 in conventional hESCs and hiPSCs. Data were obtained from^10,12,27–33^. Mean +/- SD of at least 8 biological replicates is shown.

KLF7 sustains pluripotency in conventional PSCs^27^, so we hypothesised that expression of KLF7, instead of KLF4, together with OCT4, SOX2 and cMYC (OSK7M) could enable reprogramming of primary somatic cells.

To test this hypothesis we delivered modified messenger RNAs (mmRNAs)^34,35^ encoding for OSK7M to human BJ fibroblasts, using microfluidics which has been reported to lead to rapid and efficient generation of primed iPSCs^36^. We also used OSKM and OSNL^4,6^ as positive controls. By day 14, iPSC colonies with morphology of primed pluripotent stem cells were detected (dashed circles in Fig. 2a). Immunofluorescence (IF) for pluripotency markers OCT4 and NANOG confirmed acquisition of pluripotency (Fig. 2b).

**Fig. 2:**
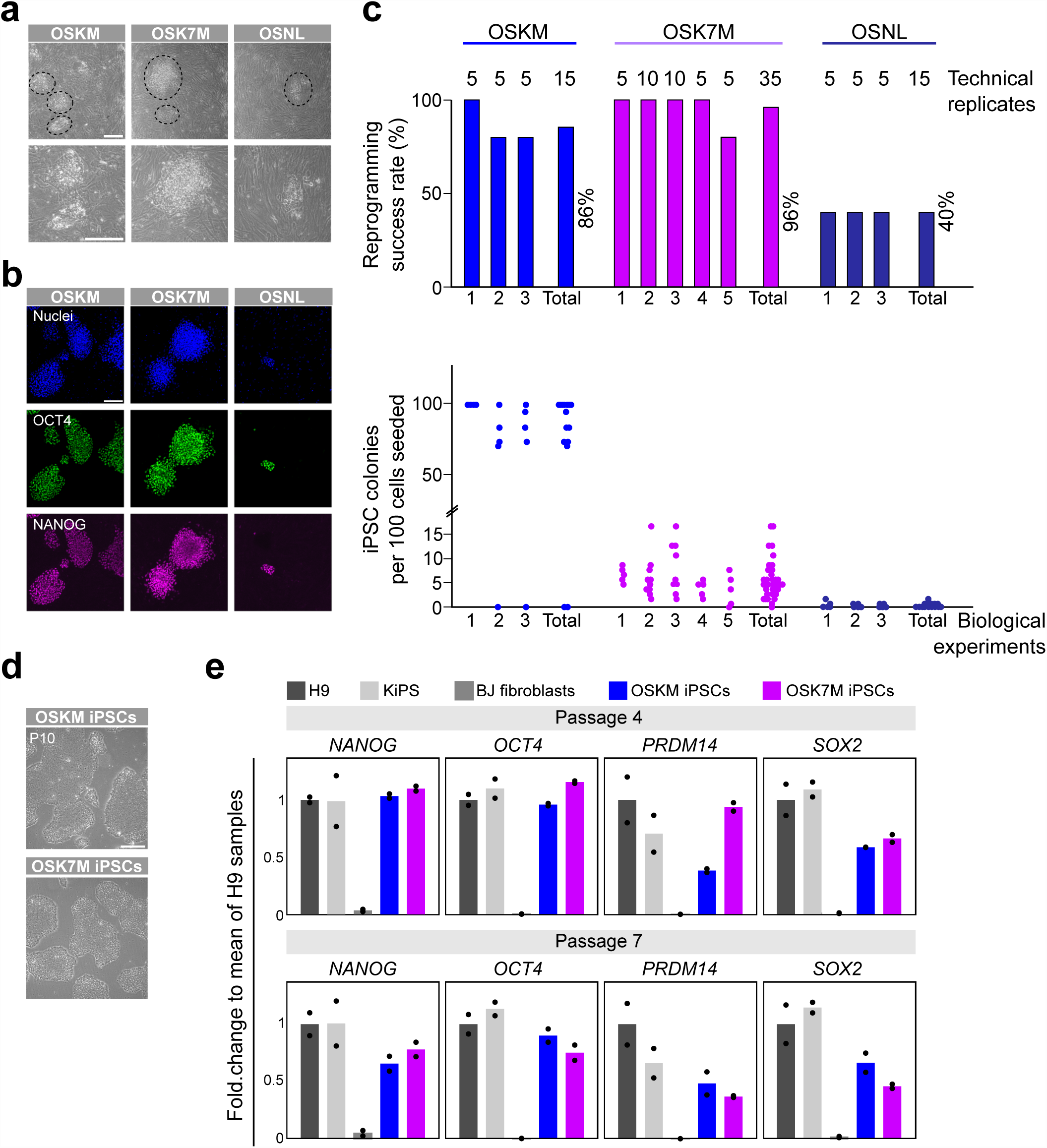
KLF7 enables somatic cell reprogramming. a: Brightfield images of iPSCs obtained at day 14 from fibroblasts reprogrammed by using 3 distinct reprogramming cocktails, OSKM, OSK7M and OSNL. Dashed circles indicate fully reprogrammed iPSCs colonies. Representative images of at least 3 independent experiments are shown. Scale bars: 100µm. b: Immunofluorescence images of pluripotency markers OCT4 and NANOG. Nuclei were stained with DAPI. Representative images of at least 3 independent experiments are shown. Scale bar: 100µm. c: Top: Reprogramming success rate calculated as percentage of technical replicates that produced at least 1 fully reprogrammed colony. Bottom: Number of iPSCs colonies obtained at day 14 from 100 cells seeded, by using 3 distinct reprogramming cocktails. Results of at least 3 independent experiments are shown. d: Representative brightfield images of stabilised iPSCs cultures (10 passages), obtained with OSKM, OSK7M and OSNL reprogramming cocktails. Representative images of at least 3 independent experiments are shown. Scale bar: 100µm. e: Barplots showing expression measured by qPCR of primed pluripotency markers in stabilised iPSCs cultures, obtained with OSKM and OSK7M reprogramming cocktails. Mean of 2 independent experiments is shown.

All three reprogramming cocktails used generated iPSC colonies, although OSNL was less robust, less efficient and formed smaller colonies (Fig. 2c and 2b). We then expanded the newly generated iPSCs to obtain stable lines. Colonies derived from OSKM and OSK7M cocktails could be readily propagated for at least 10 passages while primary iPSC colonies generated with OSNL could not stabilise in culture, therefore were not further characterised (Fig. 2d). Gene expression analysis (Fig. 2e) revealed that OSK7M iPSCs after 4 and 7 passages express pluripotency markers at levels comparable to conventional human Embryonic Stem cells (hESCs -H9) and iPSCs generated with OSKM from keratinocytes (KiPS).

These results indicate that expression of KLF7, together with OCT4, SOX2 and cMYC, enables efficient and robust human somatic cell reprogramming.

In order to further characterise these newly established iPSC lines, we performed transcriptome analyses. Unsupervised clustering showed that the transcriptional profile is clearly distinct from that of fibroblasts and comparable to hESCs (H9 cells) (Fig. 3a). Moreover, analysis of a large panel of pluripotency and fibroblasts-specific genes (Fig. 3b) further corroborated the evidence that stable iPSC lines obtained from reprogramming with OSK7M are transcriptionally indistinguishable from embryo-derived PSCs.

**Fig. 3:**
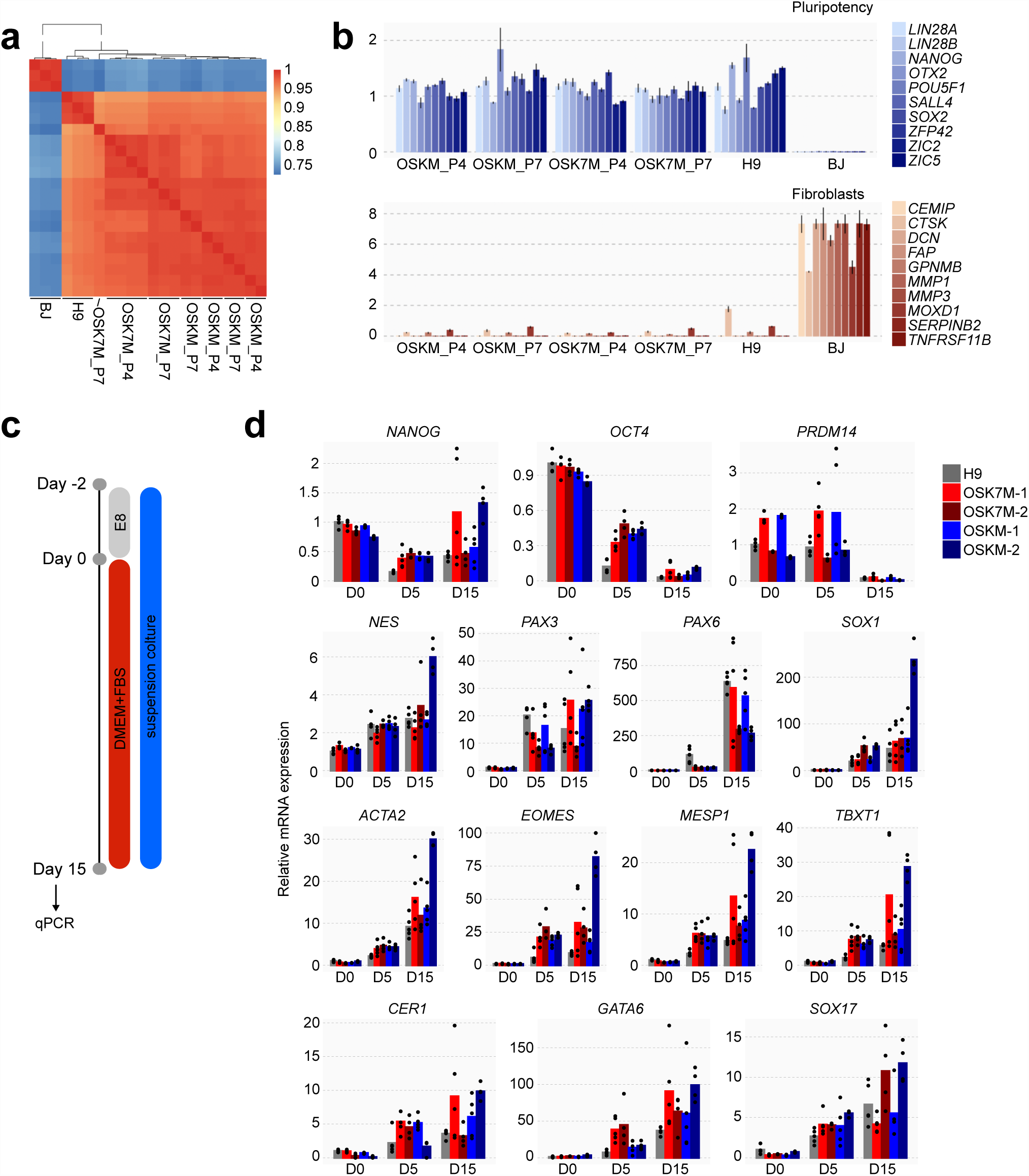
Reprogramming with OSK7M cocktail generates bone fide human iPSCs. a: Heatmap of unsupervised clustering based on transcriptome analysis in primed PSCs (H9), fibroblasts (BJ) and iPSCs obtained from reprogramming of fibroblasts with OSKM and OSK7M and stabilised for 4 and 7 passages. b: Barplots showing the mean-normalised expression of pluripotency and fibroblasts markers in iPSCs stabilised for 4 and 7 passages after reprogramming with OSKM or OSK7M. H9 PSCs and fibroblasts (BJ) were used as controls. Mean +/- SD of at least 3 biological replicates is shown. c: Schematic representation of experimental strategy used for Embryoid Bodies (EBs) differentiation of iPSCs obtained from reprogramming of fibroblasts with OSKM and OSK7M. d: Barplots showing relative mRNA expression measured by qPCR of pluripotency and lineage markers in Embryoid bodies (EBs) obtained from differentiation of iPSCs generated by reprogramming with OSKM and OSK7M cocktails. H9 PSCs were used as positive control. At least 4 biological replicates of 2 independent experiments are shown.

To assess the capacity of OSK7M iPSCs to differentiate towards the three germ layers, we performed embryoid body (EB) differentiation (Fig. 3c). After 15 days we detected reduction of pluripotency markers and comparable levels of lineage markers in all iPSC lines analysed (Fig. 3d), confirming that they are pluripotent and retain multi-lineage potential. Taken together these results clearly show that reprogramming with OSK7M generated bone fide human iPSCs.

### KLF7 improves efficiency of chemical resetting

We analysed the transcriptome of naive and conventional hPSCs and human fibroblasts, and observed that KLF7 is expressed in naive hPSCs and in primed hPSCs at comparable levels, like other known general pluripotency markers (e.g. *NANOG*, *POU5F1*, *SALL4*, *LIN28B*) (Fig. 4a and 4b).

**Fig. 4:**
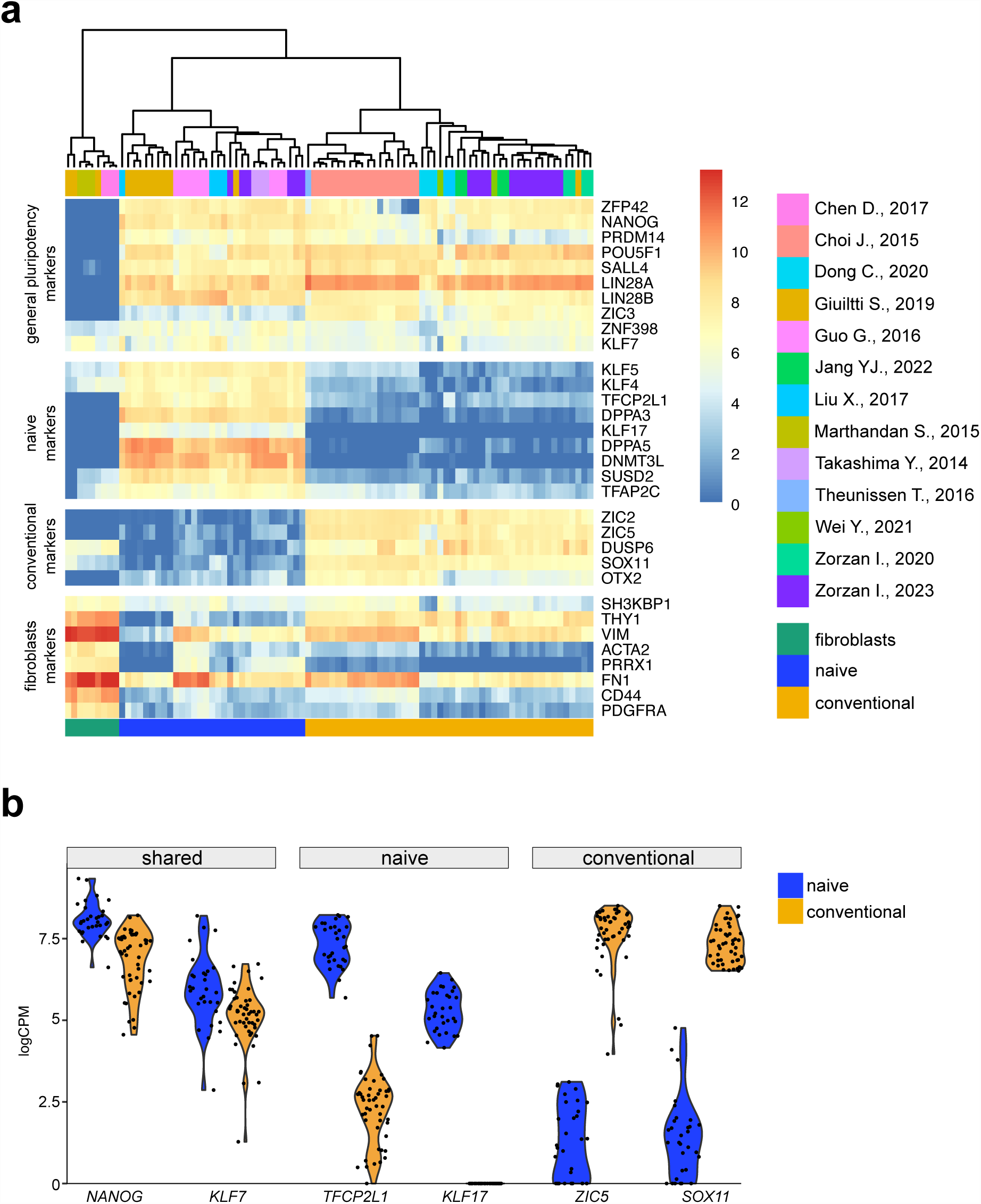
KLF7 expression in human PSCs. a: Heatmap showing expression, measured by RNAseq, of markers for naive hPSCs, conventional hPSCs, general pluripotency and fibroblasts in naive and conventional hPSCs and fibroblast cells. Data were obtained from indicated publications. b: Violin plot showing expression, measured by RNAseq, of NANOG, KLF7, TFCP2L1, KLF17, ZIC5 and SOX1 in naive and conventional hPSCs. Data were obtained from publications indicated in Fig, 4a.

In murine PSCs, Nanog and Oct4 are pluripotency factors expressed both at naive and primed state. Their forced expression, in combination with medium supporting naive pluripotency, has been shown to efficiently reset primed murine PSCs to the naive state^37,38^.

Given that KLF7 is expressed in both naive and conventional hPSCs, we hypothesised that its forced expression might boost resetting of human PSCs. Human naive PSCs can be obtained *in vitro* by chemical resetting of conventional hPSCs via transient inhibition of histone deacetylase^17^, although with low efficiency. We generated conventional iPSCs stably expressing either KLF7 (KLF7-iPSCs) or an empty vector (EMPTY-iPSCs) and then applied the chemical resetting protocol (Fig. 5a).

**Fig. 5:**
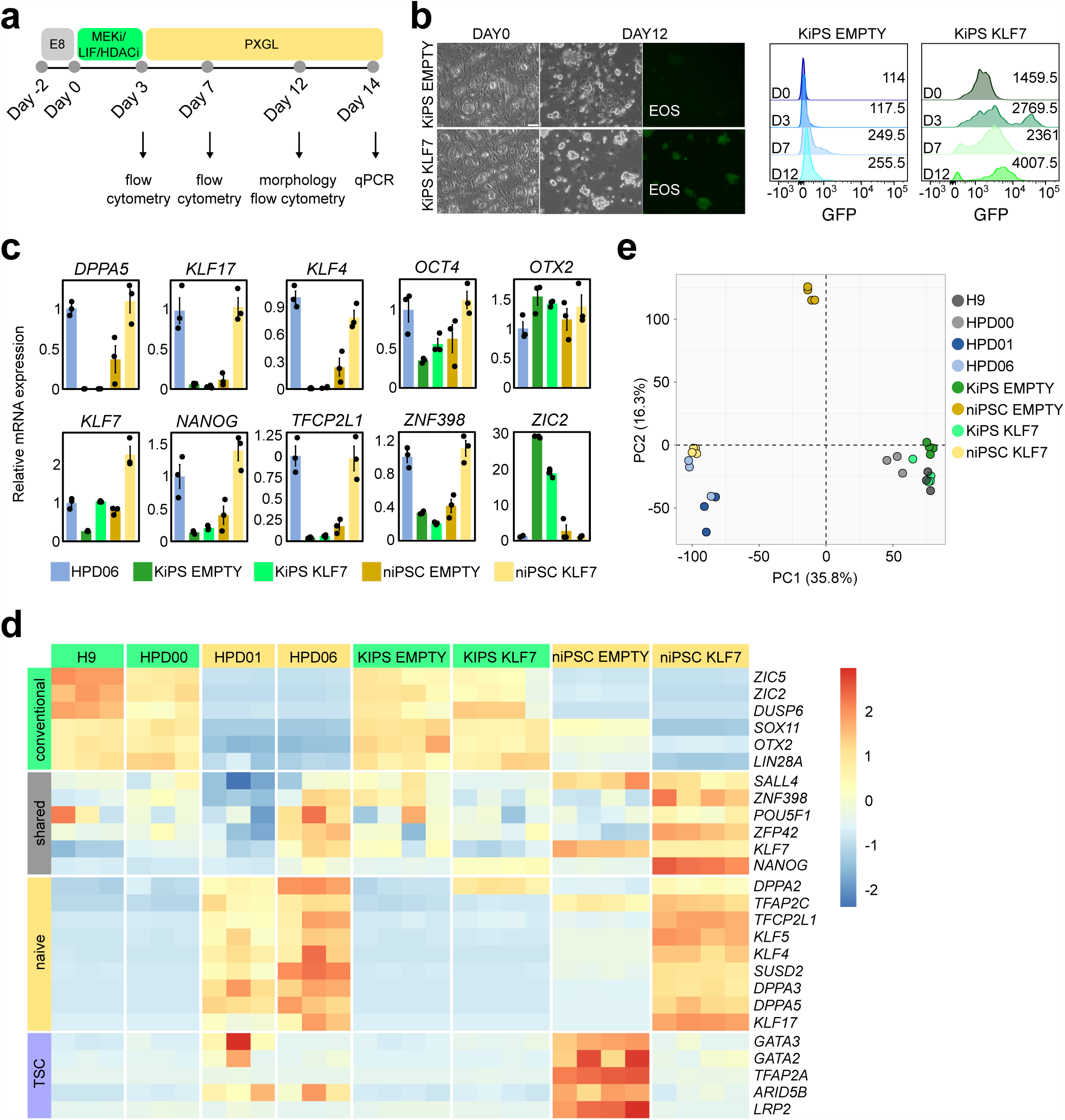
KLF7 boosts efficiency of chemical resetting. a: Schematic representation of the experimental strategy used to chemically reset conventional iPSCs to naive PSCs. b: Left: Representative images of conventional hiPSCs overexpressing an empty vector (KiPS EMPTY) or the KLF7 transgene (KiPS KLF7) at DAY0 and after 12 days of chemical resetting. Fluorescence images of OCT4-SOX2-GFP reporter EOS are shown. Representative images of 3 independent experiments are shown. Scale bar: 100µm. Right: Representative flow-cytometry plots of EOS-GFP signal in KiPS EMPTY and KiPS KLF7 cell lines at DAY0 and after 3, 7 and 12 days of chemical resetting. Representative plots of 2 independent experiments are shown. Average values of median EOS-GFP intensity of 2 independent experiments is indicated in the right part of each panel. c: Barplots showing expression measured by qPCR of naive and primed pluripotency markers in niPSC EMPTY and niPSC KLF7 obtained from chemical resetting of KiPS EMPTY and KiPS KLF7 cell lines. Naive iPSCs (HPD06) and KiPS KLF7 and KiPS EMPTY cell lines are used as controls. Mean +/- SD of 3 independent experiments is shown. d: Heatmap of Primed, Shared and Naive pluripotency genes of samples shown in Fig. 5e. Z-Scores of row-scaled expression values (CPM) are shown. e: Principal Component Analysis obtained from analysis of RNAseq data of niPSC EMPTY and niPSC KLF7 cells. At least 3 independent biological replicates were analysed.

After 7 days naive compact colonies emerged and were expanded. In EMPTY-iPSCs, resetting gave rise to a mixed population, as previously reported^17,33^, while overexpression of KLF7 resulted in a morphologically homogeneous population of naive colonies (Fig. 5b, left).

We generated EMPTY and KLF7 iPSCs stably expressing the EOS reporter, in which GFP is expressed under the control of OCT4-SOX2 responsive elements^39^, thus serving as a proxy for pluripotency. Overexpression of KLF7 led to a strong and rapid activation of EOS reporter during chemical resetting, as compared to Empty-iPSCs (Fig. 5b).

Transcriptional analysis by qPCR showed increased expression of naive markers in KLF7-naive iPSCs (KLF7-niPSCs) compared to EMPTY-niPSCs, at levels comparable to established niPSCs HPD06^10^, while the primed marker ZIC2 was inactivated in both lines (Fig. 5c). We extended our analysis to the whole transcriptome by performing RNA sequencing.

We recently reported that chemical resetting generates a mixed population of pluripotent and trophoblast cells^33^. EMPTY-niPSCs expressed high levels of trophoblast markers, retained low expression of conventional pluripotency markers (e.g. SOX11 and ZIC2) and failed to fully activate naive genes. In stark contrast, KLF7-niPSCs showed full activation of a naive transcriptional program with barely detectable expression of conventional pluripotency and trophoblast markers (Fig. 5d).

A global overview by Principal Component Analysis (PCA) revealed a similar global expression profile of KLF7 niPSCs compared to two naive iPSC lines (HPD01 and HPD06^10^) and distinct from conventional PSCs (H9, HPD00, KiPS EMPTY, KIPS KLF7) (Fig. 5e).

Taken together, our findings further endorse the observation that expression of KLF7 increases efficiency of chemical resetting, boosting expression of naive pluripotency genes while preventing activation of trophoblast markers.

### KLF7 promotes acquisition of naive identity

We then asked how, mechanistically, KLF7 promotes efficient resetting. We reasoned that it could either activate naive genes and/or block the expression of extraembryonic lineages (i.e. trophoblast). Previous transcriptional bulk analyses do not allow to discriminate whether expression of specific markers coexist in the same cells and a single-cell analysis of naive and trophoblast cells is needed. To do so, we applied the chemical resetting protocol and after 3 days of HDAC inhibition, MEK inhibition and LIF stimulation, we exposed cells to either PXGL medium to induce naive pluripotency or Trophoblast Stem Cells (TSC) medium to drive cells towards the extraembryonic fate^33^ (Fig. 6a-b). We performed quantitative IF staining for the general pluripotency marker OCT4^40–42^, the trophoblast marker GATA3^43^ and the naive PSCs marker KLF17^26^, to monitor cell fate transitions. Conventional PSCs at Day 0 (EMPTY DAY0) express OCT4 in 100% of cells and progressively acquire naive or trophoblast identity by day 12 (30% of KLF17+/OCT4+ cells at DAY12 in PXGL medium and 75.5% of GATA3+/KLF17-cells at DAY12 in TSC medium). Of note, OCT4 expression is maintained in PXGL medium (80.5% at DAY12) and completely abolished in TSC medium (2% at DAY12) (Fig. 6c-d). Interestingly cells overexpressing KLF7 display more robust activation of the naive marker KLF17 (89% of KLF17+/OCT4+ cells at DAY12 in PXGL medium) and a reduced formation GATA3+/KLF17-cells in TSC medium (44.5% at DAY12). OCT4 expression was maintained in the majority of cells exposed to PXGL (86.5% at DAY7 and 89.5% at DAY12) and, surprisingly, also in cells exposed to TSC medium (94.5% at DAY7 and 12% at DAY12).

**Fig. 6:**
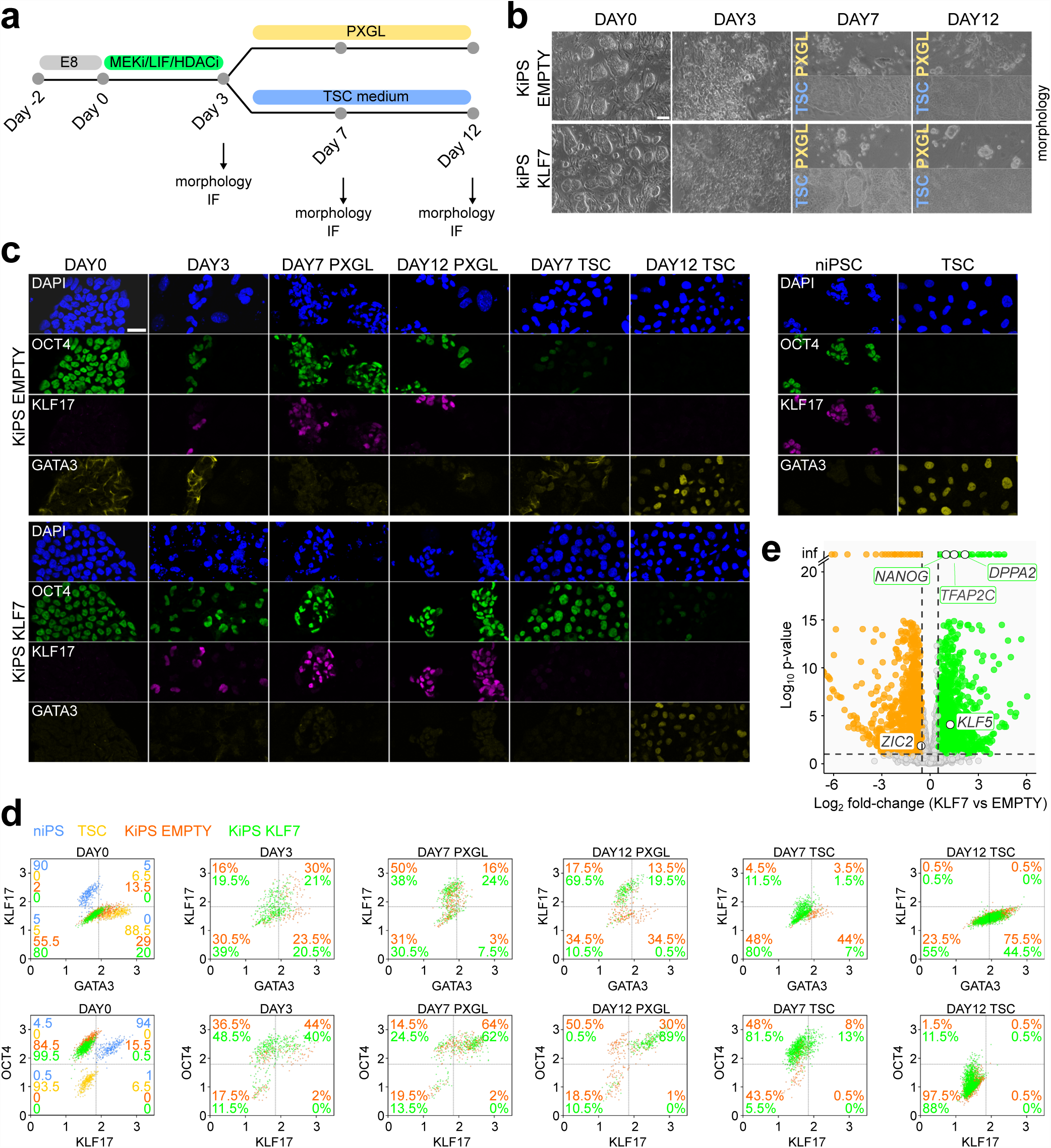
KLF7 promotes acquisition of naive identity. a: Schematic representation of the experimental strategy used for chemical resetting of KiPS EMPTY or KiPS KLF7 cells to naive or Trophoblast Stem cells (TSC), by using PXGL or TSC medium. b: Brightfield images of conventional KiPS EMPTY and KiPS KLF7 cells at DAY0 and after 3, 7 and 12 days of chemical resetting in PXGL or TSC medium, as shown in Fig. 6a. Representative images of 3 independent experiments are shown. Scale bar: 100µm. c: Immunofluorescence images of staining for OCT4, KLF17 and GATA3 in KiPS EMPTY and KiPS KLF7 cells at DAY0 and after 3, 7 and 12 days of chemical resetting in PXGL or TSC medium, as shown in Fig. 6a. niPSC and TSC were used as positive controls. Nuclei were stained with DAPI. Representative images of 2 independent experiments are shown. Scale bar: 30µm. d: Scatter plots of quantification of immunofluorescence signal for OCT4, KLF17 and GATA3. Log10 Integrated Intensity signal obtained from Cell Profile software analysis is displayed (See Materials and Methods). Representative panels of 2 independent experiments are shown. Percentage of cells in each quarter, obtained from the mean of 2 independent experiments is shown. e: Volcano plot showing transcriptome analysis of KiPS KLF7 cells compared to KiPS EMPTY cells, in standard culture conditions (E8). DOWNregulated (Log2 fold-change < -0.5 and p-value < 0.05) and UP-regulated (Log2 fold-change > 0.5 and p-value < 0.05) genes are indicated in green and orange, respectively. Known naive pluripotency markers (green) and conventional hPSCs markers (orange) are highlighted. P-values were calculated with Wald Test.

To further investigate if the mechanism of action of KLF7 in promoting pluripotency genes and blocking GATA3 is direct or indirect, we analysed the transcriptome of conventional PSCs over-expressing KLF7, as several trophoblast markers are already expressed in conventional PSCs^33^ (e.g. GATA2/3, KRT7 and TFAP2A), thus, if KLF7 is a direct repressor of trophoblast markers, we should detect a reduction in their expression in conventional PSCs. However, we observed induction of naive genes (e.g. KLF5, DPPA2, TFAP2C) and of NANOG, repression of the conventional markers ZIC2, but no significant changes in trophoblast markers (Figure 6e). Of note, TFAP2C has been shown to be required for maintenance and induction of naive pluripotency^44^, KLF5 for maintenance of naive pluripotency^24^, and NANOG for both maintenance and induction of pluripotency^15^.

Altogether these results indicate that elevated expression of KLF7 promotes the expression of naive pluripotency genes involved in maintenance and induction of pluripotency.

## DISCUSSION

In the present study, we investigated the role of KLF7 in the induction of pluripotency. We showed that KLF7 is expressed in both conventional and naive PSCs and when overexpressed in conventional PSCs it enhances chemical resetting to naive PSCs by sustaining OCT4 expression, suggesting that it rewires the core pluripotency network in maintaining the robustness of the pluripotent state.

Chemical resetting generates an initial plastic state in which naive- and TSC- specific markers are co-expressed, allowing the acquisition of pluripotent or extraembryonic fates depending on the environmental signals to which cells are later exposed^33^. We showed that KLF7 overexpression during chemical resetting promotes acquisition of the naive fate, in spite of the extraembryonic lineage. We reported that KLF7 drives acquisition of pluripotent identity also during somatic cell reprogramming.

Kruppel-like factors have been extensively studied in naive mESCs. Klf2, Klf4 and Klf5 have been characterised for their role in reprogramming of somatic and primed cells^19–21^ and in maintenance of pluripotency^18^, where only the combinatorial depletion of the three KLF factors appears to be detrimental in the maintenance of naive pluripotency and their overlapping functions target multiple naive-specific transcription factors. On the contrary, in mouse primed cells, the function of KLF factors is dispensable for their self-renewal. Transcriptional analyses of early mouse embryos confirmed expression of Klf2/4/5 only in naive pluripotent cells of the ICM or pre-implantation epiblast. Thus, in murine pluripotent cells, KLFs appear to be active in the naive state, although future studies would be needed to identify KLFs functionally relevant for murine primed pluripotency (Figure 7).

**Fig. 7:**
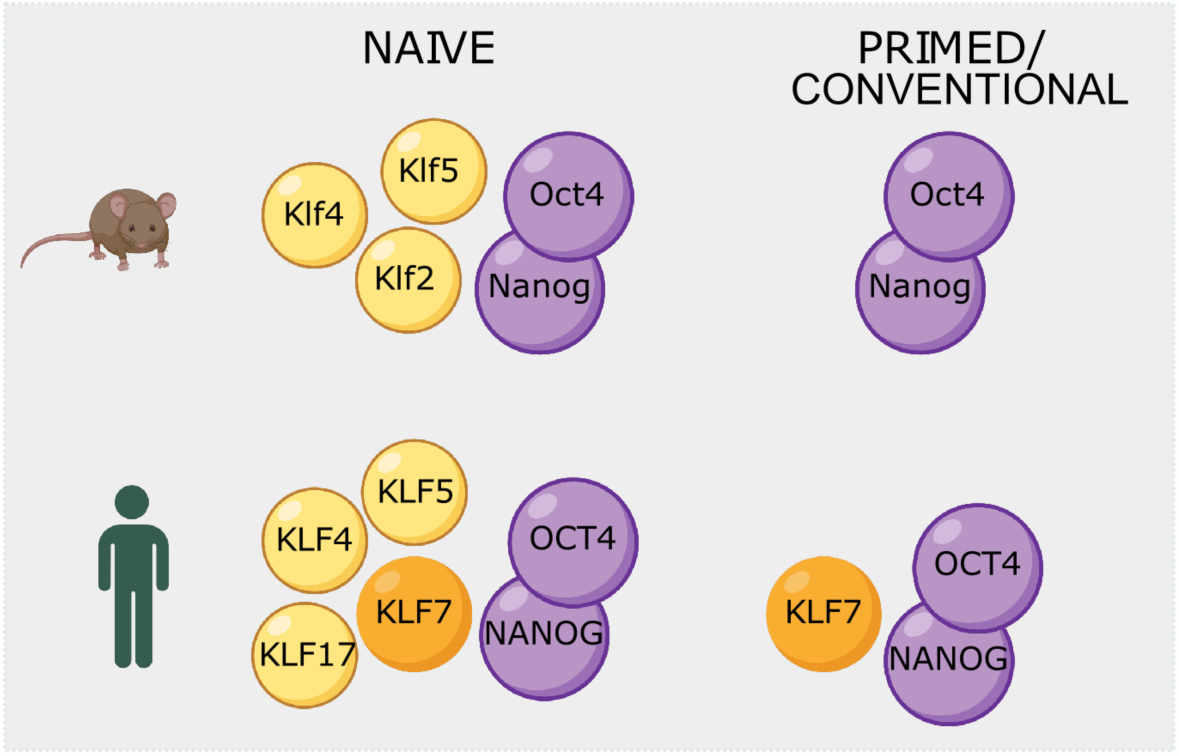
Graphical representation of Kruppel-like factors expression in naive and primed human and mouse pluripotent stem cells.

In human pluripotent stem cells, KLF4, KLF5, KLF7 and KLF17 are expressed in the naive Epiblast and in *in vitro* cultured naive PSCs^9,10,12,14,22,33^, and act as inducers of naive pluripotency.

Interestingly, KLF7 is expressed also in conventional PSCs and it promotes both their maintenance^27^ and their generation from somatic cells via reprogramming (Figure 2). Thus, in human PSCs, a single KLF acts as a general regulator of pluripotency, similarly to NANOG. NANOG is indeed expressed both in naive and primed pluripotent cells, both *in vivo* and *in vitro*, and it promotes both somatic cell reprogramming and resetting. Of note, Klf7 is not expressed at significant levels in murine PSCs, and its forced expression in primed EpiSCs is inconsequential^27^, indicating that KLF7 is a human specific pluripotency regulator, whose function is not conserved in rodents (Figure 7).

Recently, a stem cell zoo of conventional PSCs of 6 different mammalian species has been generated^45^. It would be interesting to employ those pluripotent cell lines to analyse the evolutionary conservation of KLF factors in the maintenance of pluripotency and in resetting to the naive state.

## MATERIALS AND METHODS

### Culture of hPSCs

Human primed hiPSCs (KiPS^14^, Keratinocytes induced Pluripotent Stem Cells, KiPS Empty and KiPS KLF7^27^) and HPD00^10^ and hESCs (H9) were cultured in Feeder-free on pre-coated plates with 0.5% growth factor-reduced Matrigel (CORNING 356231) (vol/vol in PBS with MgCl_2_/CaCl_2_, Sigma-Aldrich D8662) in E8 medium (made in-house according to Chen et al., 2011) at 37°C, 5% CO_2_, 5% O_2_. Cells were passaged every 3-4 days at a split ratio of 1:8 following dissociation with 0.5 mM EDTA (Invitrogen AM99260G) in PBS without MgCl2/CaCl2 (Sigma-Aldrich D8662), pH8. KiPS line was derived by reprogramming of human keratinocytes (Invitrogen) with Sendai viruses encoding for OSKM and kindly provided by Austin Smith’s laboratory.

Human naive iPSCs (HPD06 and HPD01, previously generated by direct reprogramming of somatic cells and described in Giulitti et al., 2019) were cultured on mitotically inactivated mouse embryonic fibroblasts (MEFs; DR4 ATCC) in PXGL medium^47^ or in RSeT medium (Stem Cell Technologies 05969), at 37°C, 5% CO_2_, 5% O_2_. PXGL medium was prepared as follows: N2B27 (DMEM/F12 [Gibco 11320-074], and Neurobasal in 1:1 ratio [Gibco 21103-049], with 1:200 N2 Supplement [Gibco 17502-048], and 1:100 B27 Supplement [Gibco 17504-044], 2 mM L-glutamine [Gibco 25030-024], 0.1 mM 2-mercaptoethanol [Sigma-Aldrich M3148]) supplemented with 1µM PD0325901 (Axon Medchem), 2 μM XAV939 (Axon Medchem), 2 μM Gö6983 (Axon Medchem) and 10 ng/ml human LIF (Qkine). Human naive PSCs were passaged as single cells every 4 days at split ratio 1:3 or 1:4 following dissociation with TrypLE (Gibco 12563-029) for 10 minutes at room temperature (RT). ROCK inhibitor (Y27632, Axon Medchem 1683) was added in the naive medium only for 24h after passaging. All cell lines were mycoplasma negative (Mycoalert, Lonza).

hTSCs were cultured as previously described (Okae et al, 2018). Briefly, cells were cultured on mitotically inactivated mouse embryonic fibroblasts, previously seeded on plastic plates coated with 1% gelatin, at 37°C in 5% CO_2_ and 5% O_2_. TS medium was prepared as follows: DMEM/F12 supplemented with 0.1 mM 2-mercaptoethanol, 0.2% FBS, 0.5% Penicillin–Streptomycin, 0.3%, BSA [Gibco 15260-037], 1% ITS-X [Gibco, 51500], 1.5 μg/ml L-ascorbic acid [Sigma A4544], 50 ng/ml EGF [peprotech AF-100-15], 2 μM CHIR99021 [Axon Medchem cat. nos 1386 and 1408], 0.5 μM A83-01 [Axon Medchem 1421], 1 μM SB431542 [Axon Medchem 1661], 0.8 mM VPA [HDACi, Sigma, P4543], and 5 μM Y-27632. Media was changed every 2 days and cells were passages using TrypLE Express every 3-4 days at a ratio of 1:8.

### Reprogramming

All reprogramming experiments were performed in microfluidics as previously described^35^ in hypoxia conditions (37°C, 5% CO_2_, 5% O_2_). Microfluidic channels were coated with 25 μg ml−1 Vitronectin (ThermoFisher, A14700) for 1 hour at room temperature (RT) and fibroblasts were seeded at day 0 at 15 cells per mm^2^ in DMEM/10% FBS. On day 1, 9 hours before the first mRNAs transfection, E6 medium made in-house according to Chen et al., 2011, including FGF2 100ng ml_−1_ (QKINE Cat. no Qk002 recombinant zebrafish FGF2), 1% KSR (Gibco, 10828028), ROCKi 5μM, LSD1i 0.1μM (RN-1, EMD Millipore Cat. no 489479) and 200ng ml_-1_ B18R (Invitrogen 34-8185-81) was applied. The B18R protein was added to the medium to reduce the interferon response. Cells were transfected daily at 6PM and fresh medium was given daily at 9AM. The transfection mix was prepared according to the

StemMACS_TM_ mRNA Transfection Kit (Miltenyi Biotec, 130-104-463) and using modified mRNAs (coding cocktail components) and NM-microRNAs (Stemgent StemRNA-NM Reprogramming Kit). Individual modified mRNAs (OCT4, SOX2, NANOG, LIN28A, KLF7, KLF4, MYC) were made in-house by *in vitro* transcription using mRNA synthesis with HiScribe^TM^ T7 ARCA mRNA Kit (NEB E2060S) according to the manufacturer’s instructions. During the 8 days of transfection, the dose of mRNAs was gradually increased according to cell proliferation rate and transfection-induced cell mortality. From day 9, cells were cultured in mTeSR (StemCell Technologies 05850) for 6 days to allow stabilisation of hiPSC colonies. Fresh medium was given twice a day at 9AM and 6PM.

### Embryoid bodies differentiation

For Embryoid bodies (EBs) differentiation assay, cells were detached as clumps with EDTA and plated on ultra low attachment surface plates (CORNING 3473) in E8 medium with 10 µM ROCKi. After 2 days, E8 medium was substituted with DMEM, 20% FBS, 2 mM L-glutamine, 1% NEAA and 0.1 mM 2-mercaptoethanol. Medium was changed every 2 days.

### Chemical resetting

For the chemical resetting from conventional to naive PSC or TSC, KiPS were seeded at 10000 cells/cm^2^ on mitotically inactivated MEFs in E8 medium with 10µM ROCKi (added only for 24 hours). Two days after plating (day 0), medium was changed to PD03/LIF/HDACi [N2B27 with 1µM PD0325901, 10ng/ml human LIF and 1mM VPA (HDACi)]. Following 3 days in PD03/LIF/HDACi, medium was changed to PXGL or TSC medium. Cells were passaged with TrypLE Express when confluent (around Day 8-9) on MEFs.

### Immunofluorescence

Immunofluorescence was performed on 1% Matrigel-coated glass coverslip in wells or in situ in microfluidic channels with the same protocol. For chemical resetting experiments, cells were seeded on MEFs plated on 0.5% Matrigel-coated glass coverslips at least 1 day before. Cells were fixed in 4% Formaldehyde (Sigma-Aldrich 78775) in PBS for 10 min at RT, washed in PBS, permeabilized for 1 hour in PBS + 0.3% Triton X-100 (PBST) at RT, and blocked in PBST + 5% of Horse serum (ThermoFisher 16050-122) for 5 hours at RT. Cells were incubated overnight at 4°C with primary antibodies (See Supplementary Table 2) in PBST + 3% of Horse serum. After washing with PBS, cells were incubated with secondary antibodies (Alexa, Life Technologies) (Table 1) for 45 min at RT in the case of Matrigel-coated glass coverslip and for 2 hours for staining in microfluidic devices. Nuclei were stained with either DAPI (4′,6-diamidino-2-phenylindole, Sigma-Aldrich F6057) for glass coverslip or Hoechst 33342 (ThermoFisher 62249) for microfluidic channels. Images were acquired with a Zeiss LSN700, Leica SP8 or Leica SP5 confocal microscopes using ZEN 201 or Leica TCS SP5 LAS AF (v2.7.3.9723) softwares respectively.

For chemical resetting, fluorescence intensity was quantified with Cell Profile Software (v4.1.3). DAPI staining was used to identify individual cell nuclei and the cytoplasm surrounding each nucleus. Between 4 and 7 independent fields for each sample were quantified. Integrated intensity values were plotted as log (Integrated Intensity+1). Protein presence in the cell population was determined by setting the integrated intensity threshold to 95% of the cumulative density function in samples where the protein is expressed (GATA3 in TSCs, KLF17 in niPS and OCT4 in niPS and KiPS). Thresholds used to determine percentage of positive and negative cells for a given protein were as follows: thr^GATA3^ = 1.895, thr^KLF17^ = 1.8, thr^OCT4^ = 1.74.

### Quantitative PCR

Total RNA was isolated using Total RNA Purification Kit (Norgen Biotek 37500), and complementary DNA (cDNA) was made from 500 ng using M-MLV Reverse Transcriptase (Invitrogen 28025-013) and dN6 primers. For real-time PCR, SYBR Green Master mix (Bioline BIO-94020) was used. Primers are detailed in Supplementary Table 3. Three technical replicates were carried out for all quantitative PCR. GAPDH was used as an endogenous control to normalise expression. qPCR data were acquired with QuantStudio™ 6&7 Flex Software 1.0.

### RNA sequencing and analyses

Quant Seq 3’ mRNA-seq Library Prep kit (Lexogen) was used for library construction. Library quantification was performed by fluorometer (Qubit) and bioanalyzer (Agilent). Sequencing was performed on NextSeq500 ILLUMINA instruments to produce 5 million reads (75bp SE) for the sample. For the analysis of TSCs markers, the reads were trimmed using BBDuk (BBMap v37.87), with parameters indicated in the Lexogen data analysis protocol. After trimming, reads were aligned to the Homo sapiens genome (GRCm38.p13) using STAR (v2.7.6a). The gene expression levels were quantified using featureCounts (v2.0.1). Genes were sorted removing those that had a total number of counts below 10 in at least 3 samples. All RNA-seq analyses were carried out in the R environment (v4.0.0) with Bioconductor (v3.7). We computed differential expression analysis using the DESeq2 R package (v1.28.1)52. Transcripts with absolute value of log2[FC] > 1 and an adjusted pvalue < 0.05 (Benjamini– Hochberg adjustment) were considered significant and defined as differentially expressed for the comparison in the analysis. Heatmaps were made using counts-permillion (CPM) values with the pheatmap function from the pheatmap R package (v1.0.12) on differentially expressed genes or selected markers. Volcano plots were computed with log2[FC] and −log10[adjusted p-value] from DESeq2 differential expression analysis output using the ggscatter function from the ggpubr R package (v0.4.0). Barplots were made using CPM values with the ggbarplot function from the ggpubr R package. For the time point analysis during resetting and the following stabilisation in PXGL or TSC medium, transcript quantification was performed from raw reads using Salmon (v1.6.0)53 on transcripts defined in Ensembl 105. Gene expression levels were estimated with tximport R package (v1.20.0)54 and differential expression analysis was computed using the DESeq2 R package (v1.28.1)52. Transcripts with absolute value of log2[FC] ≥ 3 and an adjusted p-value < 0.05 (Benjamini–Hochberg adjustment) were considered significant. Principal component analysis was performed on variance stabilised data (vst function from DESeq2 R package v1.32.052) using prcomp function on the top 5000 most variable genes. Heatmaps were performed using the pheatmap function (pheatmap R package v1.0.12) on log2 count per million (CPM) data of selected markers. All analysis except salmon were performed in R version 4.1.1.

### Analyses of published RNA sequencing data were performed as follows

Raw-reads were downloaded for samples from relative databases as described in Supplementary Table 1. Transcript quantification was performed using Salmon (v1.9.0)^48^ with transcripts defined in Ensembl 106. Gene expression levels were estimated with tximport R package (v1.26.1)^49^. Batch correction was performed using ComBat_seq function from sva R package (v3.46.0; https://bioconductor.org/packages/release/bioc/html/sva.html). Batches have been defined following library preparation: batch 1 for full length libraries and batch 2 for Quant Seq 3′ mRNA-seq Library Prep kit of the GEO series GSE184562.

CPM on batch corrected counts were computed using the cpm function of edgeR package (v3.40.2;^50^). PCA was performed using the svd r function on log transformed CPM.

All plots except heatmap have been done using ggplot2 v3.4.2 (https://ggplot2.tidyverse.org/authors.html#citation). Heatmap was done using pheatmap R package (v1.0.12). All analyses were performed using R v4.2.2.

### Flow cytometry

After dissociation in single-cell suspension using TrypLE, cells were resuspended in PBS and filtered. Cells were analysed according to EOS-GFP expression with BD FACSCantoTM II cytometer and BD FACSDivaTM (v. 9.0) software. Data analysis was performed with FlowJo software (v10.9.0).

### Statistics and reproducibility

For each dataset, sample size n refers to the number of independent experiments or biological replicates, shown as dots, as stated in the figure legends. R software (v4.0.0) was used for statistical analysis. Error bars indicate standard deviation or the standard error of the mean (SEM), as stated in the figure legends.

## Acknowledgements

G.M.’s laboratory is supported by grants from the Giovanni Armenise–Harvard Foundation, the Telethon Foundation (GJC21157), Microsoft Research and ERC Starting Grant (MetEpiStem).

## Author contributions

GM conceived the project. MA, IZ, EC designed the experiments and interpreted the results. MA and IZ performed reprogramming experiments and molecular analyses. IZ and EC performed chemical resetting experiments and molecular analyses. MP performed mmRNA synthesis for reprogramming experiments. MA and PM performed bioinformatics analyses. EC and GM wrote the paper, with help from all authors.

## Data availability

Sequencing data that support the findings of this study have been deposited in the Gene Expression Omnibus (GEO). Previously published RNAseq data that were re-analysed here are available under accession codes listed in Supplementary Table 1. Primers and antibodies used are listed in Supplementary Tables 2 and 3. Source data are provided with this study. All other data supporting the findings of this study are available from the corresponding authors on reasonable request.

